# Rho-dependent termination and RNase E-mediated cleavage: Dual pathways for RNA 3’ end processing in polycistronic mRNA

**DOI:** 10.1101/2024.10.01.616006

**Authors:** Heung Jin Jeon, Monford Paul Abishek N, Xun Wang, Heon M. Lim

**Author notes:** To whom correspondence may be addressed. **E-mail:** Heung Jin Jeon & Heon M. Lim. These authors contributed equally to this work (Heung Jin Jeon, Monford Paul Abishek N).

## Abstract

“Pre-full-length” transcripts are produced at the end of the polycistronic galactose (*gal)* operon, 5’*galE-galT-galK-galM3’*, via Rho-dependent (RDT) and -independent transcription termination (RIT). The full-length *galETKM* mRNA’s 3’ end is acquired by exo-nucleolytic processing of the 3’-OH ends of the pre-full-length transcripts. However, the *gal* operon produces an mRNA named *galE* whose 3’ end forms at ∼120 nucleotides from the *galE* stop codon, thus in the following gene, *galT*, establishing polarity in gene expression. In this study, we investigated the molecular processes that generate the 3’ end of *galE* mRNA. We discovered that the 3’ ends of pre-galE mRNA are produced in the middle of the *galT* gene as a result of the combination of two separate molecular processes - one previously reported as RDT and the other as unreported RNase E-mediated transcript cleavage. The 3’ ends of the pre-*galE* mRNA are exo-nucleolytically processed to the current 3’ end of the *galE* mRNA. A hairpin structure of 8 base-pair stems and 4 nucleotide-loop formed 5-10 nucleotides upstream of the 3’ ends of the *galE* mRNA blocks the exoribonuclease digestion and renders stability. These findings showed that RNase E produces RNA 3’end establishing polarity in gene expression, in contrast to the general role of mRNA degradation.

**Significance statement:** Here, we show the findings of two molecular mechanisms that generate the pre-*galE* mRNA 3’ends in the *gal* operon: Rho-dependent termination (RDT) and RNase E-mediated cleavage. These 3’ ends are subsequently processed to produce stable *galE* mRNA with a hairpin structure that prevents exoribonuclease degradation. This mechanism establishes gene expression polarity by generating the 3’ end of *galE* mRNA within the *galT* gene, contrasting with the usual mRNA degradation role of RNase E. The study reveals a unique role of RNase E in mRNA processing and stability.

## Introduction

In *Escherichia coli*, transcription initiated from the promoter terminates by the two transcription termination mechanisms; Rho-dependent (RDT) and Rho-independent (RIT) (Reviewed in (1)). In principle, the termination process reveals the 3’ end of the transcript, the final nucleotide containing a free 3’OH group (1). The 3’ end of an emerging transcript formed through transcription termination does not signify the 3’ end of mature RNA (1). The nascent RNA’s 3’ end undergoes immediate exoribonuclease digestion (3’→5’), where a hairpin structure formed near the 3’ end of the nascent RNA blocks the exoribonuclease digestion and renders stability to the mature mRNA (2, 3).

Transcription initiated from the *P1* and *P2* promoters of the *gal* operon terminates at the end of the operon and generates the full-length mRNA referred to as *galETKM*, which harbors the open reading frames of the 4 structural genes of the operon, *galE, galT, galK*, and *galM* (Fig. 1A) (4, 5). The terminator hairpin functions as the RIT terminator and blocks the 3’ to 5’ exoribonuclease digestion by RDT, making the stable 3’ end of the *galETKM* mRNA (2). Transcriptions that go through the terminator hairpin were terminated at the Rho-dependent terminator; Rho-terminated transcripts have been quickly processed by exonuclease digestion (2, 6). Thus, transcription terminates at the RIT or RDT in the *gal* operon (2, 7).

**Figure. 1.**
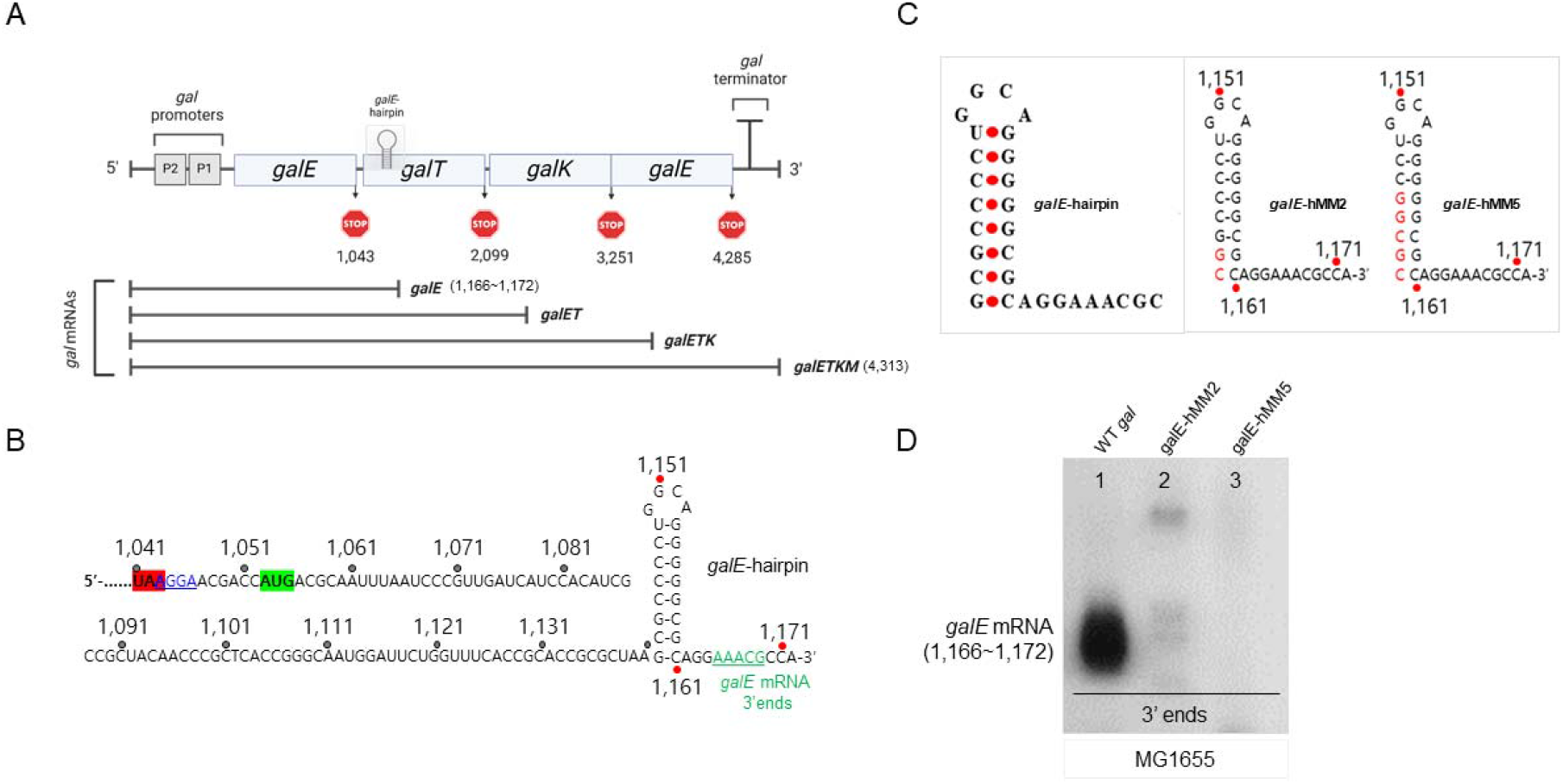
A hairpin structure, *galE*-hairpin is responsible for the 3’ end of *galE* mRNA functioning as an exo-block. A) galactose operon. Numbers indicate the nucleotide residue coordinate of the *gal* operon, which starts from the transcription initiation site of the *galP1* promoter. Northern blot E DNA probes (500 bp) were created via Polymerase chain reaction (PCR) amplification with primers corresponding to the *galE* region (from +27 to +527 in *gal* coordinates), subsequently radiolabeled with 32^P^ as previously described. B) Nucleotide sequence of the 3′ end of the *galE* mRNA (green) in the WT *gal* operon. The *galE* stop codon is highlighted in red and the *galT* initiator codon and SD sequence are highlighted in green and blue (underlined), respectively. The *galE*-hairpin structure is depicted based on base complementarity between positions 1,142 and 1,161. C) *galE*-hMM2 and *galE*-hMM5 mutants. The *gal* mutants were generated in the single-copy plasmid, pGal where the entire *gal* operon is cloned and assayed in MG1655 cells from where the entire *gal* has been removed, MG1655Δ*gal*. D) 3’RACE of *galE* mRNA 3’ends of WT *galE*-hMM2 and *galE*-hMM5 mutants.

The *gal* operon produces not only the full-length mRNA but also 3 additional mRNA species, *galETK, galET*, and *galE*, having 3’ ends at the end of each gene, *galK, galT*, and *galE*, respectively (Fig. 1A) (5, 7-9). The sRNA, Spot 42 binding at the intercistronic sequence of *galT-galK* of mRNA causes RDT and RNase E-mediated transcript cleavage (10-12).

Production of these *gal* mRNA species, *per se*, establishes polarity in gene expression, and higher gene expression in the promoter-proximal genes than in the distal genes (5, 13, 14). Thus, the cause of the gene expression’s polarity is deeply rooted in the molecular processes to generate the 3’ ends of these *gal* mRNA species (5, 6, 12). In this study, we show that at the 3’ end of the smallest mRNA species from the *gal* operon, the *galE* mRNA is generated from two different mechanisms: RDT (previously reported) (6) and an unreported RNase E-mediated transcript cleavage. We found that RNase E-mediated transcript RNA cleavage also generates the 3’ end of the *galE* mRNA. These results indicated that, in contrast to the typical role of RNase E in degrading mRNA, RNase E-mediated transcript cleavage might result in the formation of the mRNA’s 3’ end (10). This suggests that transcription termination may not be the sole method for creating the 3’ end of mRNA and that RNase E cleavage could be employed to control polarity in the *gal* operon.

## Results

### The 3’ end of *galE* mRNA is processed from those of two ‘pre-*galE*’ mRNA which is blocked by a hairpin structure

The *galE* mRNA appears to be about 1.2 kilobases (kb) in a northern blot when probed with the E-probe that hybridizes to the first half of the *galE* gene (5). The 3’ end of *galE* mRNA is generated at 1,166-1,171 (*gal* coordinate starts from the first nucleotide of the *P1* transcript) about 130 nucleotides downstream from the stop codon of *galE*, and about 120 nucleotides downstream from the initiator codon of *galT* (Fig. 1A) (6). RNA secondary structure analysis suggested that at the 3’ ends of *galE* mRNA, a stem with 6 consecutive G: C and 1 U: G base-pairings and a loop with 4 nucleotides could be formed at 5-10 nucleotides upstream from the major 3’ ends of the of *galE* mRNA (1,166-1,172) (Fig. 1B) (6).

Nucleotide changes in the *gal* operon DNA were done in a single-copy plasmid, pGal where the entire *gal* operon is cloned (*SI Text*). We assayed, the resulting *gal* mutants in MG1655 cells from where the entire *gal* has been removed, MG1655Δ*gal* (8). We removed two base pairs from the bottom of the stem of the *galE*-hairpin by changing the 1,161 cytidines and 1,160 guanines to their complementary nucleotide; guanine and cytidine, respectively (Fig. 1B and C). The resulting *gal* mutant, gal-*EHMM2*, failed to produce the 3’ ends at 1,166-1,172 (lane 2 in Fig. 1C). We further removed 5 base pairs from the stem by changing the nucleotides from 1,161 to 1,156 to their complementary nucleotides (Fig. 1B). The resulting mutant, gal-*EHMM5* also failed to produce the 3’ end at 1,166-1,172 (lane 3 in Fig. 1D). These results demonstrated that the *galE*-hairpin is a critical factor for the generation of the 3’ end of the *galE* mRNA at 1,166-1,172. The role of the *galE*-hairpin likely is to block exoribonuclease digestion that is initiated somewhere downstream and renders the stability to the mature mRNA (6).

To locate the genetic *loci* where the exoribonuclease digestion initiates molecular processes that cause the 3’ ends of *galE* mRNA, we inserted an additional *galE*-hairpin at 1,200 and created a *gal* mutant, double hairpin at 1,200 (*DH1200*), where the second hairpin starts at 1,200 and ends at 1,219 (Fig. 2A) (6). The 3’RACE results of the *DH1200* mutant showed that the 3’ ends of *galE* mRNA decreased ∼50±2% of WT (Fig. S1) (6). Interestingly, we observed that the ∼50±2% decrease in the 3’ ends of *galE* mRNA was caused by the fact that the 3’ ends at 1,169-1,172 (Upper 3’ends) are not produced (Fig. S1). Nevertheless, in the *DH1200* mutant, we did observe a new cluster of 3’ ends at 1,224-1,230, 5-11 nucleotides downstream of the foot of the stem of the inserted *galE-*hairpin (Fig. S1) (6). These results suggest that the second hairpin reduced *galE* mRNA 3’ ends by half of the 3’ ends of *galE* mRNA, possibly by blocking 3’ to 5’ exonuclease digestion downstream of position 1,200 where the hairpin was inserted. In the *DH1200* mutant, there is an absence of production of the upper 3’ ends at position 1,169-1,172, while there is a newly observed production of the 3’ ends at position 1,219-1,223. This suggests that the 3’ ends of *galE* mRNA are generated through RNA processing from two separate pre-*galE* mRNAs, each having 3’ ends located at different positions; 1) pre-*galE1*_;_ having 3’ ends between 1,171-1,200, and 2) pre-*galE2*; having 3’ ends downstream of 1,200.

**Figure. 2.**
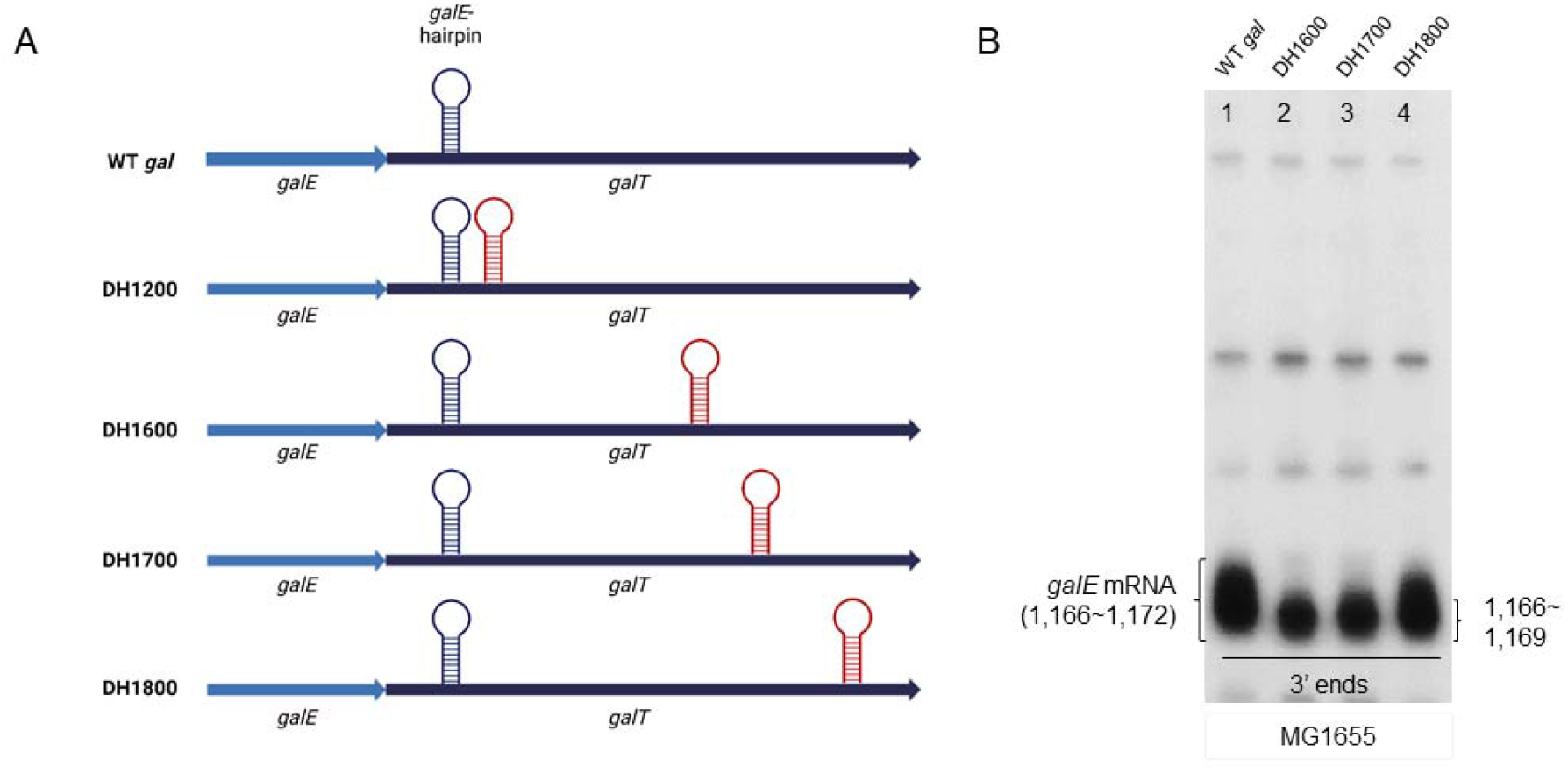
Exoribonuclease digestion initiates molecular processes that cause the 3’ ends of *galE* mRNA. A) Schematic of Double hairpin (DH) mutants; *DH1200, DH1600, DH1700*, and *DH1800*. B) 3’RACE of *galE* mRNA of WT and *DH1200-1800* mutants.

To locate the 3’ end of pre-*galE2*, we inserted the *galE*-hairpin at 1,600, 1,700, and 1,800 positions, generating *DH1600, DH1700*, and *DH1800* (Fig. 2A). We assayed the 3’ end of the *galE* mRNA at 1,166-1,172 in these mutants. When the second *galE*-hairpin is located 400 nucleotides downstream of 1200 (*DH1600*), the 3’ end of the *galE* mRNA decreased with an absence of production of the upper 3’ ends at position 1,169-1,172. When the second *galE*-hairpin is located 500 and 600 nucleotides downstream of 1200 (*DH1700 and DH1800)*, we notice a gradual increase in the 3’ end of the *galE* mRNA at 1,166-1,172. Its formation reached 55±3% of WT in *DH1600*, and 65±4% of WT in *DH1700*. It increased to as much as WT in *DH1800* (lanes 2, 3, and 4, Fig. 2B). These findings suggested that inserting a hairpin every 100 nucleotides into the system exposed RNase E cleavage sites, which improved RNA stability and accumulation over time by blocking 3’ to 5’ exo-nucleolytic digestion blocked by the inserted-hairpins, resulting in a gradual increase in *galE* mRNA levels. The 3’RACE assay does not detect these 3’ ends, because these 3’ ends are likely subjected to 3’ to 5’ exo-nucleolytic digestion which is blocked by the *galE-*hairpin to make the 3’ end of the *galE* mRNA.

### RNase E-mediated transcript cleavage and RNase II processing of terminated RNA generates the 3’ end of pre-*galE2*

Our previous study reported that RDT occurs at the end of the *galE* gene in the *gal* operon (6). The Rho-inhibitor bicyclomycin (BCM) experiments demonstrate that RDT generates the 3’ end of pre-*galE1* (1,183) (6). We hypothesized that the 3’ end of pre-*galE2* could be generated by an endo-ribonucleolytic cleavage on a *gal* transcript. Thus, we expected that the *galE* mRNA would decrease in the precise endoribonuclease mutant cells like RNase E as it’s the major endoribonuclease in *E. coli* (15-17). To determine if this nuclease is accountable for producing the 3′ ends of pre-*galE2*, we examined its existence in the temperature-sensitive GW20 (*ams1ts*) strain for RNase E activity (16). Our northern blot assay revealed that the amount of *galE* mRNA decreased by approximately 70% at the non-permissive temperature (44ºC) compared to the permissive temperature (30ºC). In contrast, no significant changes were observed in the levels of other *gal* mRNAs, except for *galETK* and *galETKM* (Fig. 3A and Fig. S2). This might be due to RNase E cleaving the polycistronic *gal* operon transcript in a way that destabilizes *galE* mRNA while stabilizing other parts, like *galETK* and *galETKM*, leading to increased levels of downstream mRNAs. The 3’ RACE assay showed that the 3’ ends of *galE* mRNA decreased ∼60±6 % of WT, importantly only the upper 3’ends at 1,169-1,172 were formed, suggesting that the 3’ end of pre-*galE2* could be generated by transcript RNA cleavage by RNase E (Fig. 3B and C).

**Figure. 3.**
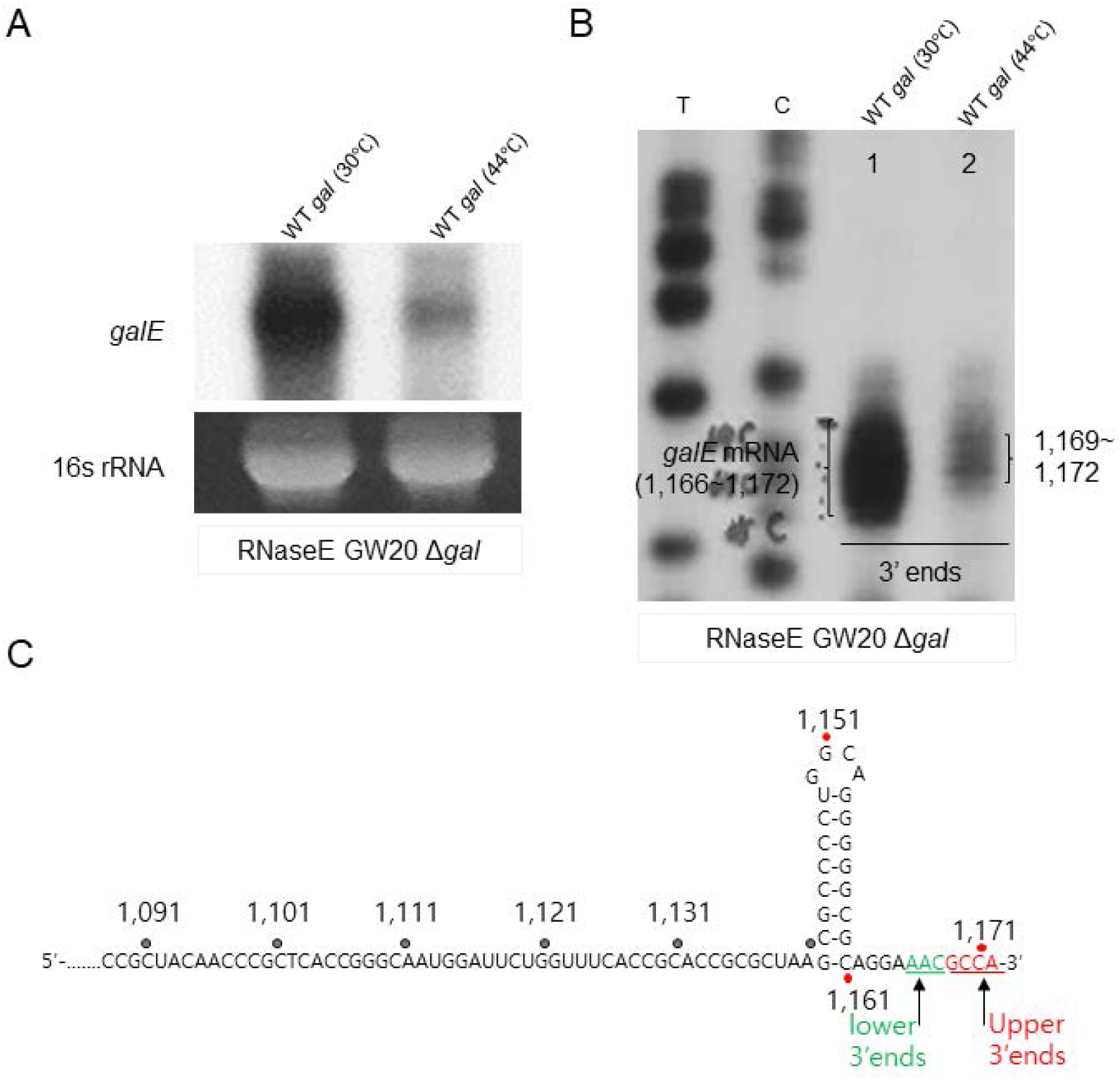
RNase E-mediated endo-nucleolytic cleavage processing is the source of the 3’ ends of pre-*galE*. A) Northern blot and B) 3’RACE assay of the *galE* mRNA in GW20Δ*gal* (temperature-sensitive RNase E mutant) cells. Cells were cultured at both permissive (30°C) and non-permissive (44°C) temperatures for analysis. C) Nucleotide sequence of the upper (red) and lower (green) 3′ ends of the *galE* mRNA based on the 3’RACE assay on B).

RNase II aids in the degradation of RDT-generated mRNA 3’end in the *trp* operon of *E. coli* and bacteriophage T3 (18). We used the Δ*gal*Δ*rnb* strain (RNase II deleted) (2) with both the WT and DH1200 variant *gal* plasmids to investigate this. Interestingly, 3′ RACE assays of the *gal* transcripts revealed that the 3′ ends at position 1,166-1,172 shifted to 1,169-1,172 (upper 3’ends) in WT *gal* (Fig. 4A), and the 3′ end at position 1,166-1,169 shifted to 1,169-1,172 in *DH1200* (inserted second hairpin) with the 3’ ends at 1,219-1,223 disappearing (lanes 3 and 4, Fig. 4B). The *DH* mutants *DH1200, DH1700*, and *DH1800* were subjected to a 3’RACE assay in GW20 (*ams1ts*), to investigate the stability and presence of *galE* mRNA 3’ ends. The results from the 3’RACE assay demonstrate that the *DH1200, DH1700*, and *DH1800* mutants in GW20 temperature-sensitive strain show a significant loss of mRNA 3’ ends at the non-permissive temperature of 44°C (Fig. 4C) and more importantly only the 3’ upper ends at 1,169-1,172 were formed. Based on these results we hypothesize, that post-RDT events downstream indicate that an unidentified RNase likely processed the Rho-terminated 3′ ends from 1,183 to 1,169-1,172, and RNase II would then have removed the final three nucleotides from the 3′ ends, changing it from 1,169-1,172 to 1,166, making the final stable 3’ends of *galE* mRNA at 1,166-1,172 (Fig. 3C). This also confirms that in *DH1200* (inserted second hairpin), 3’ ends at 1,219-1,223 could be a result of RNase E-mediated transcript cleavage where the second hairpin blocked 3’ to 5’ exonuclease digestion of the 3’ end of ‘pre-*galE2*’.

**Figure. 4.**
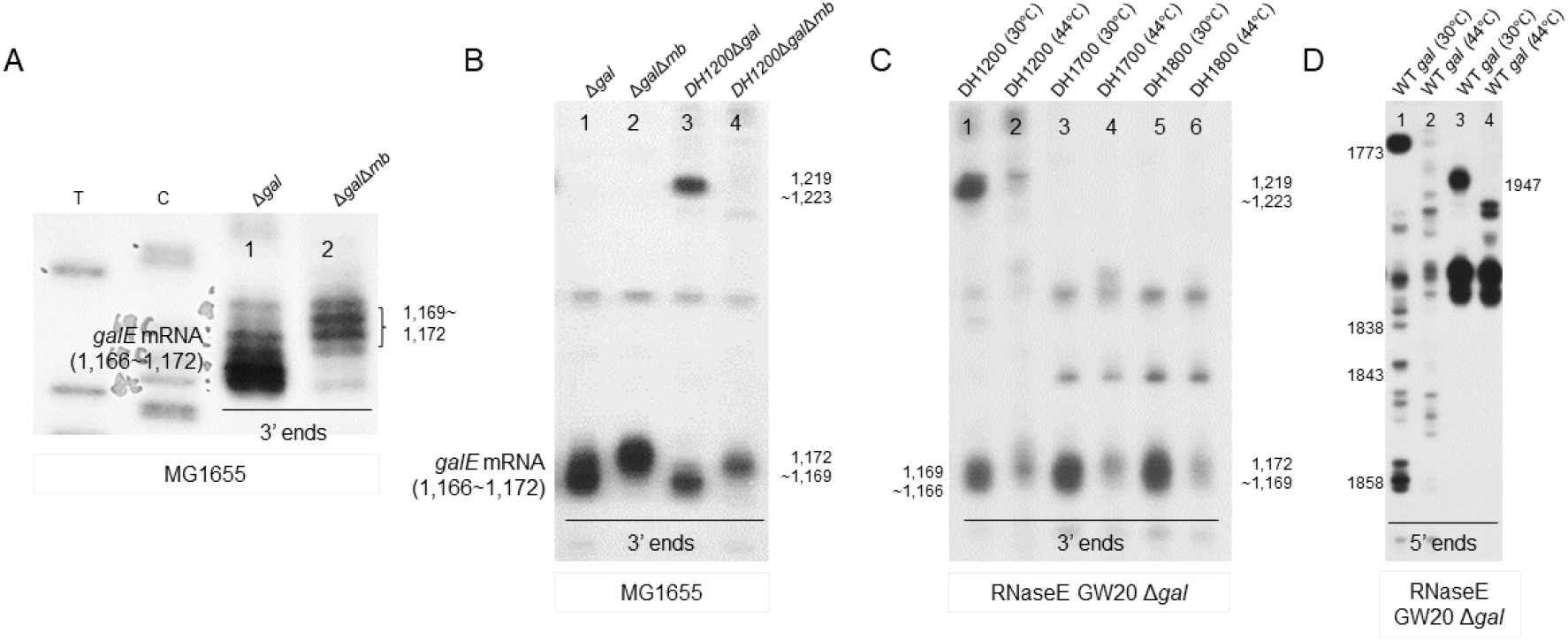
RNase II-mediated exo-nucleolytic processing contributes to the generation of the 3’ ends of pre-*galE*. A) 3’RACE assay of the *galE* mRNA in Δ*gal*Δ*rnb*, where RNaseII is deleted from the chromosome. B) 3′ RACE assay of *galE* mRNA in Δ*gal*Δ*rnb* harboring the DH1200 mutant variant. T and C: DNA sequencing ladder. C) 3’RACE assay of the *galE* mRNA in GW20Δ*gal* (temperature-sensitive RNase E mutant) cells harboring the DH mutants (DH1200, 1700 and 1800). D) 5’RACE assay to identify RNaseE cleavage in GW20Δ*gal* cells. Cells were cultured at both permissive (30 °C) and non-permissive (44 °C) temperatures for analysis.

Since RNase E cleavage can lead to changes in the abundance and integrity of RNA transcripts, understanding where and how RNase E cleaves transcripts will help decipher the post-transcriptional control mechanisms. To determine the exact location of the 3’ end of pre-*galE2*, we investigated the 5’ ends of RNA between positions 1,700 and 1,900, rather than directly searching for the 3’ ends. We performed 5’ RACE on total RNA from the GW20Δ*gal* strain carrying the pGal plasmid. At the permissive temperature (30ºC), several 5’ ends were identified through the RACE assay (Lane 1, Fig. 4C). Notably, at the non-permissive temperature (44ºC), these 5’ ends were absent (Lane 2, Fig. 4C). These findings suggest that RNase E may endonucleolytically cleave the *gal* transcript at these positions *in vivo*. Moderate sequence specificity is associated with RNase E cleavage, and it is characterized by a consensus sequence: 5′-R(A/G)NW(A/U)UU-3′ (R = A or G, N = any nt, W = A or U) (17, 19), where RNase E cuts between residues, N and W, resulting with the W residue at the 5′ end (17). DNA sequence analysis indicated, that consensus sequences of RNase E-cleavage sites were found between positions 1,700 and 1,900.

Our *in vivo* and sequence analysis demonstrate that RNase E cleaves the *gal* transcript between residues between positions 1,700 and 1,900, resulting in 5’ ends at positions (lane 2 in Fig. 4C). These findings not only support the idea that RNase E-mediated cleavage generates the 3’ end of pre-*galE2* but also reveal that aberrant cleavages by RNase E downstream 1,700 leading to a reduction in the 3’ end of *galE* mRNA at 1,166 and 1,172. By identifying the cleavage sites, the 5’ RACE assay provides insights into how RNase E influences the stability and maturation of RNA transcripts, such as the *galE* mRNA, which is critical for elucidating the mechanisms of gene regulation at the RNA level.

### RNase E-mediated transcript cleavage and RDT both regulated *galE* mRNA 3’ end *in vivo*

At the *galE-galT* junction, the stop codon of *galE* and the initiator codon of *galT* are separated by nine nucleotides. To examine the effect of removing *galT* translation initiation on transcription, we changed the initiator codon of *galT* (AUG) to an ordinary codon (AAA), generating a *gal* mutant (*galT* start*º*) for northern blotting. The northern blot of the *galE* mRNA in the *galT* start*º* showed a very thick and smeared RNA band starting at 2.0 kb (lane 3 of Fig. 5A) down to the beginning of the operon, and a diminished full-length *galETKM* and *galET* (Fig. S3A). These results show that most of the *gal* transcription in the *galT* start*º* mutant was terminated downstream and a few transcription events reached the end of the operon. Based on these data, we suggest that RDT and possibly degradation of the 3’ ends of Rho-terminated transcript mRNAs are the cause of the thick and smeared band in the *galT* start*º* mutant. We treated the *galT* start*º* mutant with BCM (Rho-inhibiting) for 10 min and found that the thick smearing band decreased (lane 4, Fig. 5A), demonstrating that, indeed, the thick smearing band was caused by RDT. These results suggest that the absence of *galT* translation initiation caused a break in the transcription-translation coupling, which in turn, caused RDT.

**Figure. 5.**
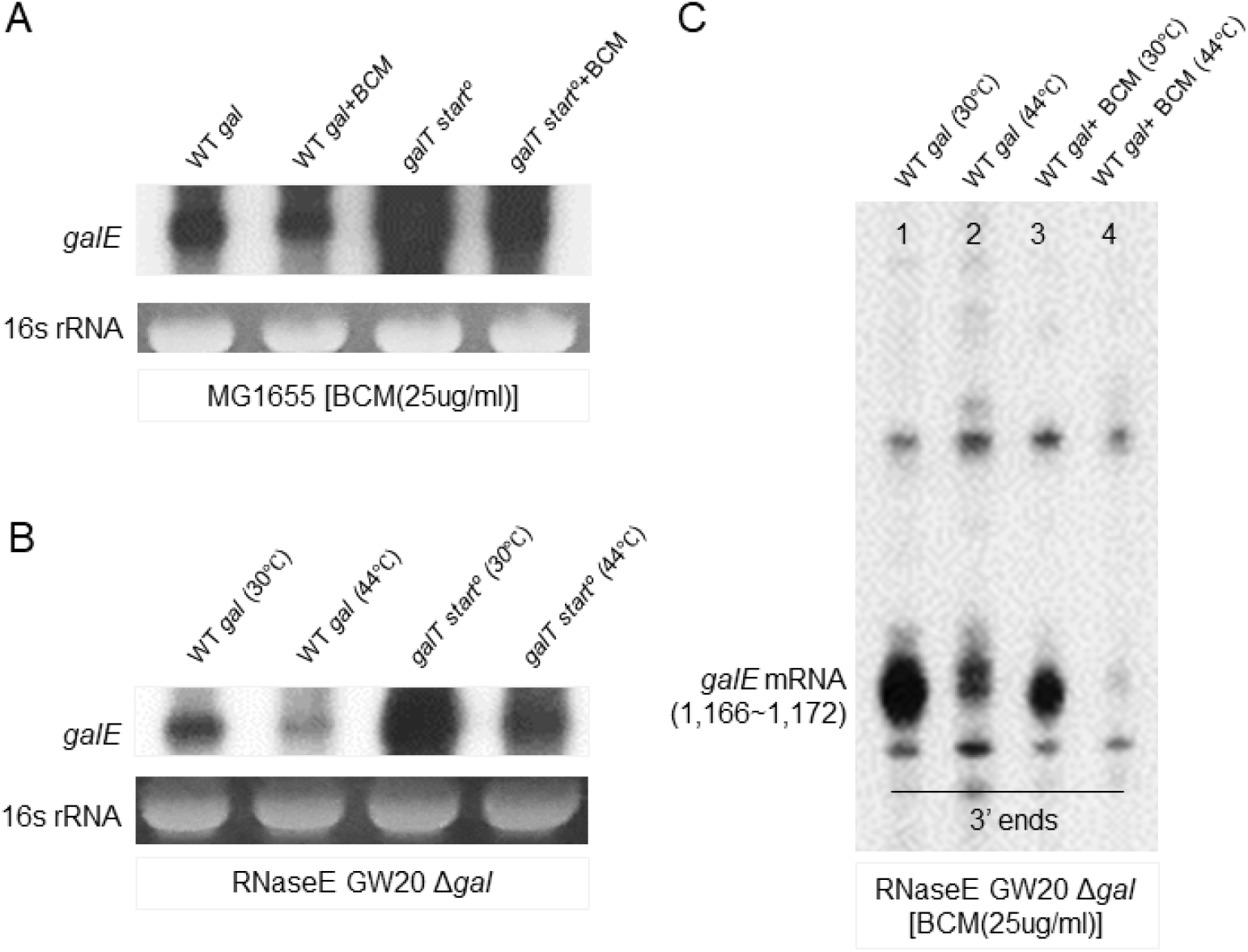
Rho and RNase E affecting *galE* mRNA. A) Northern blot analysis of *galE* mRNA in *galT* startº mutant with or without the Rho inhibitor bicyclomycin (BCM). To determine if *RDT produces galE mRNA*, we treated MG1655 cells at an OD600 of 0.6 in LB medium with 25µg/mL BCM concentration for 10 minutes. B) Northern blot analysis of *galE* mRNA in *galT* startº mutant in GW20Δ*gal*. C) 3’RACE assay of *galE* mRNA 3’ends in GW20Δ*gal* cells with or without the Rho inhibitor bicyclomycin (BCM). D) Relative band intensity calculated from the 3’RACE analysis of C).

To investigate the effect of RNase E-mediated cleavage of the *galT* start*º* mutant on *galE* mRNA expression, we performed the northern blot assay in the GW20 (*ams1ts*) strain. At the permissive temperature (30°C), a distinct band corresponding to the *galE* mRNA is observed (lane 1 in Fig. 5B). In the *galT* start*º*, at 30°C, the *galE* mRNA band appears significantly increased, indicating increased expression of *galE* mRNA in the mutant strain under permissive temperature (lane 3 in Fig. 5B and Fig. S3C). At the non-permissive temperature (44°C), the *galE* mRNA band appears significantly diminished (lane 2 in Fig. 5B). In *galT* start*º*, at 44°C, the *galE* mRNA band is reduced (lane 4 in Fig. 5B) compared to *galT* start*º*, at 30°C (lane 3 in Fig. 5B), indicating substantial repression by RNase E of *galE* mRNA (Fig. S3C). These results suggest that the absence of *galT* translation initiation could also cause RNase E-mediated transcript cleavage.

However, the source of the break-in transcription-translation coupling in the *galT* start*º* mutant is likely rooted in translation termination at the stop codon of *galE* (6, 20). We imagined that the transcription-translation break could be prevented if the translation termination of *galE* was removed from the *galT* start*º* mutant; continuous translation without interruption by termination at the stop codon of *galT* could eliminate the break in transcription-translation coupling. To test this idea, we changed the stop codon of *galE* (TAA) to an ordinary codon (AAA) in *galT* start*º*, generating a double mutant *galET-one-frame* for northern blotting. As expected we found that the thick smeared band indicative of RDT disappeared, and the *galETKM* band was restored to the level of WT (lane 4 in Fig. S3B). These results demonstrate that continuous translation activity prevented the break in transcription-translation coupling. These results were suggestive that if translation termination of *galE* is removed from WT cells, the 3’ end of the *galE* mRNA at 1,166-1,172 should decrease in amount, because the continuous translation activity at the *galE-galT* cistron junction would prevent the break in the transcription-translation coupling, thus preventing RDT from occurring.

Data from this study suggest that multiple RNase E cleavages occur between 1,700 and 1,900, and the 3’ end generated at 1,183 by RDT is processed to the galE mRNA 3’ end at positions 1,166-1,172. Thus, if we assay the 3’ end of the *galE* mRNA at 1,166-1,172 in GW20 (*ams1ts*) strain in the presence of BCM (inhibiting Rho), we expected that at the non-permissive temperature (inhibiting RNase E), the 3’ end at 1,166-1,172 would disappear. The 3’RACE assay of the 3’ ends at 1,166-1,172 in GW20 cells showed, indeed that is the case (Fig. 5C). Without BCM, GW20 cells generated about half of the 3’ ends at positions 1,166-1,172 at the non-permissive temperature (lane 2 in Fig. 5C). This supports the previous finding that RNase E is responsible for producing roughly half of the 3’ ends at these positions. When BCM was added, GW20 cells produced about half of the 3’ ends at positions 1,166-1,172 at the permissive temperature (lane 3 in Fig. 5C), reinforcing the earlier result that Rho contributes to the generation of the other half of these 3’ ends. Nonetheless, in the presence of 40 µg/ml BCM, the 3’ ends at 1,166-1,172 was hardly detected at the non-permissive temperature (lane 4 in Fig. 5C). When we assayed the 3’ end of the *galE* mRNA at 1,166-1,172 in GW20 (*ams1ts*) strain in the presence of *galE* stop° (inhibiting Rho) (6, 20), similar results were seen in the 3’RACE assay (Fig. S4A and B). This supports the idea that the 3’ ends at 1,166-1,172 in WT cells originate from two distinct sources, with both RDT and RNase E-mediated cleavage contributing equally to their formation.

## Discussion

### Translation initiation failure and regulation of mRNA 3’ ends

#### Stochastic failure of translation initiation of the leading ribosome evokes RDT

In prokaryotes, transcription and translation are closely coupled processes (21-27). Ribosomes can bind and initiate translating the nascent mRNA as soon as the RNA polymerase synthesizes it. When translation begins, the first ribosome on the mRNA is considered the leading ribosome (Fig. 6A). The nascent mRNA will not be adequately shielded by ribosomes if the leading ribosome is unable to initiate translation (6, 28). Rho protein binds to the mRNA at specific sites called Rho utilization (rut) sites (2, 29-33) (Fig. 6B). When ribosomes translate the mRNA, they hinder Rho from reaching these locations by physically obstructing the mRNA (21, 30, 34). Suppose the process of translation initiation is unsuccessful, the unoccupied mRNA is then available to Rho, which proceeds to unravel the RNA-DNA hybrid in the transcription bubble, ultimately leading to the termination of transcription (6, 35).

**Figure. 6.**
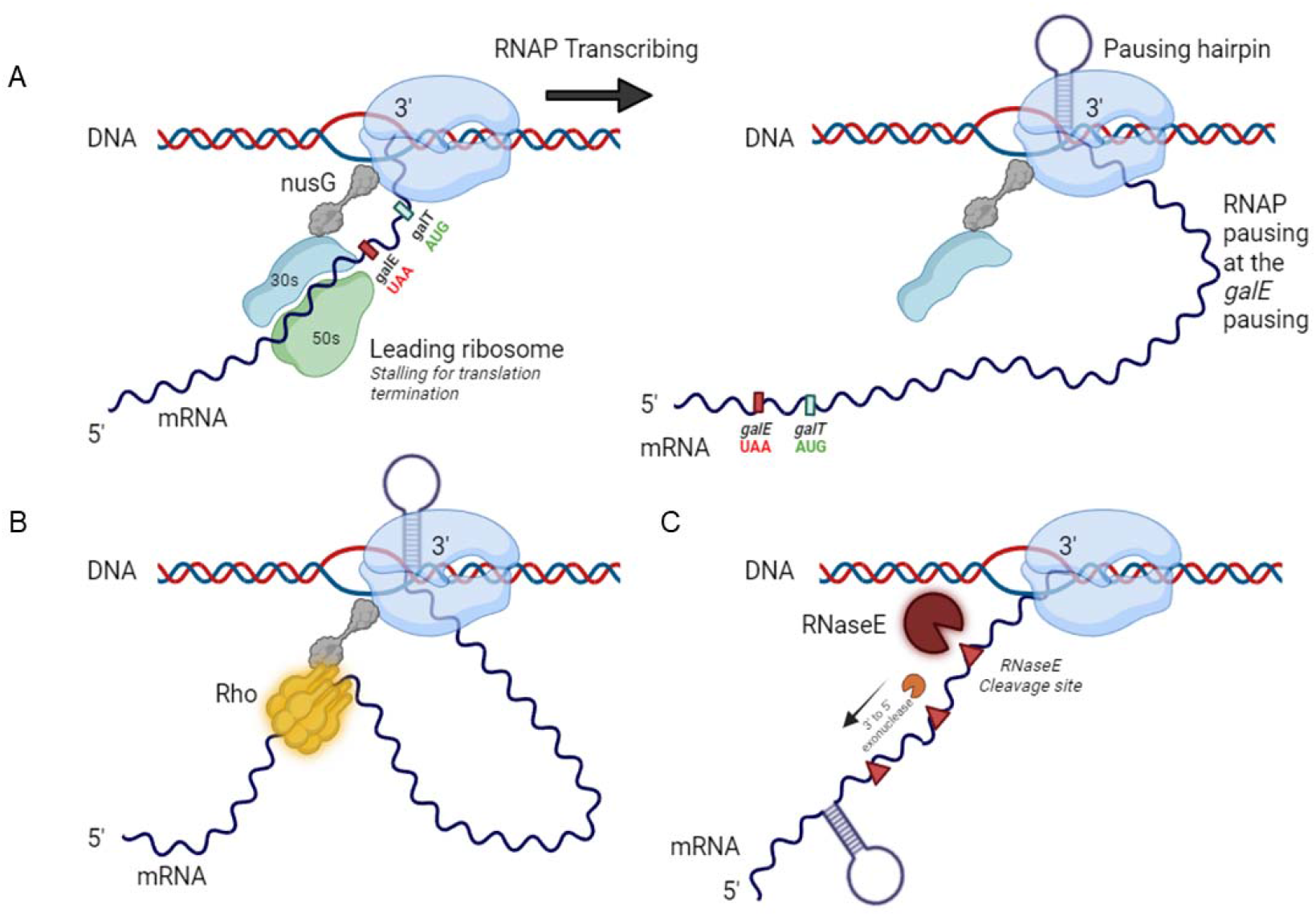
Transcription model. A) Model representing a transcription-translation complex in which the leading ribosome’s 30S ribosome remains linked to the paused RNA polymerase prolonged by *galE-*hairpin following the cessation of *galE* translation termination. B) Model of transcription termination in which the RNA polymerase is stalled downstream of the *galE* stop codon, and Rho is brought to it by the NusG protein. C) RNase E-mediated cleavage model, in which RNase E cleaves the transcript’s RNA free from the ribosome and subjected to 3’ to 5’ exo-nucleolytic digestion, which is blocked by the *galE*-hairpin to produce the 3’ end of the *galE* mRNA.

In the *gal* operon, normally, transcription is continuous, but about 10% of transcription terminates prematurely at the end of the *galE* gene due to RDT (Fig. 6A and B) (6, 7). This termination is influenced by whether the translation of the next gene, *galT*, successfully initiates. The failure to initiate translation of the *galT* gene after *galE* translation is completed leads to RDT at the *galE-galT* cistron junction (Fig. 6B) (6). The sequences of nucleotides at the junctions of cistrons might have evolved to enable random failures in the initiation of translation, which could impact the proportion of mRNA generated for various genes in an operon (6, 36).

#### Stochastic failure of translation initiation of an ordinary ribosome could cause RNase E-mediated transcript cleavage

In the second instance when translation initiation fails stochastically, in the absence of a bound ribosome, certain regions of the mRNA, particularly those near the ribosome binding site (RBS) and the start codon remain unprotected exposing the RNase E cleavage sites (37). RNase E functions as an endoribonuclease which involves triggering mRNA degradation by cleaving particular sites in the RNA molecule (3, 15, 38). When translation initiation by the ribosome does not occur, these unprotected regions become open to RNase E (37). Using Northern blot and RACE analysis, this study shows that RDT and RNase E-mediated cleavage contribute to the formation of the *galE* mRNA (Fig. 6B and C). We report that this degradation not only prevents the synthesis of the protein encoded by the mRNA but can also influence the stability of the RNA. When RNase E activity is inhibited, as seen in the temperature-sensitive strain GW20, the levels of these mRNA species drastically decrease, confirming RNase E’s role in their production (Fig. 3 and 5).

RNase E cleaves downstream of the *galE* hairpin, facilitating the decay process by allowing 3’→5’ exoribonuclease access for digestion of the remaining RNA (Fig. 1, 5, and 6C) (18). This cleavage ensures the complete degradation of the mRNA, contributing to efficient RNA processing. RNase E is involved in the synthesis of mRNA rather than the degradation of mRNA because target mRNA *galE* decreases when the cleavage leads to degradation of the target mRNA increases in amount. In the case of *galETKM* mRNA, *galETKM* increases when the RNase E mutant cells are at the non-permissive temperature. mRNA stability factors can influence natural or mutational polarity in the *gal* operon. Given the role of RNAse E, it’s plausible that its mutation could lead to changes in mRNA stability and, consequently, polarity (5). At the non-permissive temperature (44°C), the altered activity of RNAse E potentially affects its impact on polarity, suggesting that the RNAse E mutant may partially release natural or mutational polarity in the *gal* operon at this temperature (Fig. S2) (5). This research indicates that in *E. coli*, the gene downstream must start the translation process for transcription to continue beyond intercistronic junctions. If this translation initiation does not occur, it leads to the cessation of transcription, which can be facilitated by either Rho or RNase E. These results emphasize the multipart relationship between transcription and translation in the regulation of bacterial genes and offer new possibilities for understanding operon dynamics and mRNA stability.

### Regulation of mRNA 3’ end in the polycistronic *gal* operon

The generation of the 3’ end of RNA by RDT and RNase E cleavage is a prevalent mechanism in the formation of *galE* and *galETKM* mRNA within the galactose operon (Fig. 6). The *gal* operon is essential for the metabolism of galactose in *E. coli* and includes several genes that are transcribed from a shared promoter (Fig. 1A) (39-41). For *galE* mRNA, this process ensures the production of a stable and functional transcript necessary for encoding UDP-galactose 4-epimerase, an enzyme critical for galactose metabolism (42). In the case of *galETKM* mRNA, RDT and RNase E cleavage produce a polycistronic mRNA that includes the *galE* gene along with *galT, galK, and galM*, encoding enzymes that convert galactose into glucose-1-phosphate. These coordinated mechanisms of 3’ end generation are crucial for the regulation and efficient expression of the galactose operon, enabling the cell to effectively manage and utilize galactose.

## Materials and methods

### Bacterial strains, growth conditions and primers

MG1655 Δ*gal*, MG1655Δ*gal*Δ*rnb* (RNaseII deletion) (2), and GW20 (*ams1ts*; RNase E temperature-sensitive mutant) (8, 9, 20, 37) were the *E. coli* strains used in this study. The strains of *E. coli* with chromosomal deletions were created using phage Lambda Red-mediated recombineering MG1655 (43). Bicyclomycin (BCM) was a generous gift from Max E. Gottesman (Columbia University, USA). Primers used in this study are listed in the Table S1.

### Plasmids

The construction of the pHL1277 plasmid involved the insertion of the galactose operon, spanning from position -75 to +4333, between the *EcoR*I and *BamH*I sites of the pCC1BAC vector (Epicenter Biotechnologies, USA). The galactose operon was amplified from genomic DNA using PCR primers listed in Table S1. For site-directed mutagenesis, custom synthetic primers containing the specific desired mutations were designed for PCR amplification. Following the amplification of DNA fragments containing different mutants of the galactose operon by PCR from the pHL1277 plasmid, these fragments were employed as “mega primers” for the subsequent round of PCR. The resulting PCR fragments were then digested with EcoRI and HindIII and subsequently ligated into pHL1277. This process led to the generation of several derived plasmids including pHL1751 (EHMM2), pHL1754 (EHMM5), pHL1930 (DH1200), pHL1931 (DH1600), pHL1932 (DH1700), pHL1933 (DH1800), pHL1657 (*galE* stopº mutant), pHL1658 (*galT* startº mutant), and pHL1939 (*galET*-ONE-frame).

### Total RNA extraction and Northern blot analysis

Total RNA was extracted from 2 × 10^8^ *E. coli* cells using the Direct-zol™ RNA MiniPrep kit (Zymo Research) as previously described (8, 37). For the northern blot assay, the RNA samples were resolved by gel electrophoresis and transferred overnight to a positively charged nylon membrane (Ambion, USA; TurboBlotter, Whatman, UK). The nylon membranes were hybridized and washed as per the manufacturer’s recommendations (Ambion, United States) (8, 37). ImageJ software was used to calculate the relative intensity of the RNA bands (NIH).

### Rapid amplification of cDNA ends (RACE)

The RNA preparation was treated with Turbo DNase I (Thermo Fisher Scientific, USA) to eliminate any DNA contamination. The 3’RACE and 5’RACE assays were conducted according to previously described methods to amplify either the 3’ or 5’ RNA ends (44-47). ImageJ software was used to calculate the relative intensity of the RNA bands (NIH).

## Acknowledgments

This work was funded by the National Research Foundation of Korea (NRF-RS202300243742) to H.J. The authors thank Prof. Max E. Gottesman (Columbia University, USA) for his generous gift of BCM.

## Author Contributions

H.J., and H.L.: Conceptualization, funding acquisition, project administration, and supervision; H.J., and M.N.: Data analysis, data curation, investigation, validation, and visualization; H.J.: Methodology; X.W.: Validation, and visualization; M.N., H.J., and H.L.: Writing – original draft preparation, review & editing.

## Additional files

**Supplemental material** (*SI Appendix, PDF file)*

Figures S1-S4 and Table S1.

## References

1. A. Ray-Soni, M. J. Bellecourt, R. Landick, Mechanisms of Bacterial Transcription Termination: All Good Things Must End. Annu Rev Biochem 85, 319–347 (2016).

2. X. Wang et al., Processing generates 3’ ends of RNA masking transcription termination events in prokaryotes. Proc Natl Acad Sci U S A 116, 4440–4445 (2019).

3. M. P. Hui, P. L. Foley, J. G. Belasco, Messenger RNA degradation in bacterial cells. Annu Rev Genet 48, 537–559 (2014).

4. S. Adhya, Suboperonic regulatory signals. Sci STKE 2003, pe22 (2003).

5. H. J. Lee, H. J. Jeon, S. C. Ji, S. H. Yun, H. M. Lim, Establishment of an mRNA gradient depends on the promoter: an investigation of polarity in gene expression. J Mol Biol 378, 318–327 (2008).

6. H. J. Jeon, M. P. A. N Y. Lee, H. M. Lim, Failure of Translation Initiation of the Next Gene Decouples Transcription at Intercistronic Sites and the Resultant mRNA Generation. mBio 13, e0128722 (2022).

7. M. P. A. N H. Jeon, X. Wang, H. M. Lim, Reporter Gene-Based qRT-PCR Assay for Rho-Dependent Termination In Vivo. Cells 12 (2023).

8. X. Wang et al., Expression of each cistron in the gal operon can be regulated by transcription termination and generation of a galk-specific mRNA, mK2. J Bacteriol 196, 2598–2606 (2014).

9. H. J. Jeon, Y. Lee, M. P. A. N C. Kang, H. M. Lim, sRNA expedites polycistronic mRNA decay in Escherichia coli. Front Mol Biosci 10, 1097609 (2023).

10. H. J. Jeon et al., sRNA-mediated regulation of gal mRNA in E. coli: Involvement of transcript cleavage by RNase E together with Rho-dependent transcription termination. PLoS Genet 17, e1009878 (2021).

11. T. Moller, T. Franch, C. Udesen, K. Gerdes, P. Valentin-Hansen, Spot 42 RNA mediates discoordinate expression of the E. coli galactose operon. Genes Dev 16, 1696–1706 (2002).

12. X. Wang, S. C. Ji, H. J. Jeon, Y. Lee, H. M. Lim, Two-level inhibition of galK expression by Spot 42: Degradation of mRNA mK2 and enhanced transcription termination before the galK gene. Proc Natl Acad Sci U S A 112, 7581–7586 (2015).

13. S. Adhya, M. Gottesman, Control of transcription termination. Annu Rev Biochem 47, 967–996 (1978).

14. B. De Crombrugghe, S. Adhya, M. Gottesman, I. Pastan, Effect of Rho on transcription of bacterial operons. Nat New Biol 241, 260–264 (1973).

15. J. E. Clarke, L. Kime, A. D. Romero, K. J. McDowall, Direct entry by RNase E is a major pathway for the degradation and processing of RNA in Escherichia coli. Nucleic Acids Res 42, 11733–11751 (2014).

16. M. Wachi, G. Umitsuki, K. Nagai, Functional relationship between Escherichia coli RNase E and the CafA protein. Mol Gen Genet 253, 515–519 (1997).

17. J. G. Belasco, Ribonuclease E: Chopping Knife and Sculpting Tool. Mol Cell 65, 3–4 (2017).

18. J. E. Mott, J. L. Galloway, T. Platt, Maturation of Escherichia coli tryptophan operon mRNA: evidence for 3’ exonucleolytic processing after rho-dependent termination. EMBO J 4, 1887–1891 (1985).

19. Y. Chao et al., In Vivo Cleavage Map Illuminates the Central Role of RNase E in Coding and Non-coding RNA Pathways. Mol Cell 65, 39–51 (2017).

20. H. J. Jeon, N. Monford Paul Abishek, Y. Lee, J. Park, H. M. Lim, Transcription Needs Translation Initiation of the Downstream Gene to Continue Downstream at Intercistronic Junctions in E. Coli. Curr Microbiol 81, 89 (2024).

21. R. S. Washburn et al., Escherichia coli NusG Links the Lead Ribosome with the Transcription Elongation Complex. iScience 23, 101352 (2020).

22. C. Wang et al., Structural basis of transcription-translation coupling. Science 369, 1359–1365 (2020).

23. S. Proshkin, A. R. Rahmouni, A. Mironov, E. Nudler, Cooperation between translating ribosomes and RNA polymerase in transcription elongation. Science 328, 504–508 (2010).

24. B. M. Burmann et al., A NusE:NusG complex links transcription and translation. Science 328, 501–504 (2010).

25. M. W. Webster et al., Structural basis of transcription-translation coupling and collision in bacteria. Science 369, 1355–1359 (2020).

26. V. Svetlov, E. Nudler, Unfolding the bridge between transcription and translation. Cell 150, 243–245 (2012).

27. G. M. Blaha, J. T. Wade, Transcription-Translation Coupling in Bacteria. Annu Rev Genet 56, 187–205 (2022).

28. N. Said et al., Steps toward translocation-independent RNA polymerase inactivation by terminator ATPase rho. Science 371 (2021).

29. L. V. Richardson, J. P. Richardson, Rho-dependent termination of transcription is governed primarily by the upstream Rho utilization (rut) sequences of a terminator. J Biol Chem 271, 21597–21603 (1996).

30. E. Skordalakes, J. M. Berger, Structure of the Rho transcription terminator: mechanism of mRNA recognition and helicase loading. Cell 114, 135–146 (2003).

31. V. Epshtein, D. Dutta, J. Wade, E. Nudler, An allosteric mechanism of Rho-dependent transcription termination. Nature 463, 245–249 (2010).

32. D. Dar, R. Sorek, High-resolution RNA 3’-ends mapping of bacterial Rho-dependent transcripts. Nucleic Acids Res 46, 6797–6805 (2018).

33. Z. Hao, V. Svetlov, E. Nudler, Rho-dependent transcription termination: a revisionist view. Transcription 12, 171–181 (2021).

34. Y. Murayama et al., Structural basis of the transcription termination factor Rho engagement with transcribing RNA polymerase from Thermus thermophilus. Sci Adv 9, eade7093 (2023).

35. V. Molodtsov, C. Wang, E. Firlar, J. T. Kaelber, R. H. Ebright, Structural basis of Rho-dependent transcription termination. Nature 614, 367–374 (2023).

36. E. S. Komarova et al., Influence of the spacer region between the Shine-Dalgarno box and the start codon for fine-tuning of the translation efficiency in Escherichia coli. Microb Biotechnol 13, 1254–1261 (2020).

37. H. J. Jeon et al., Translation Initiation Control of RNase E-Mediated Decay of Polycistronic gal mRNA. Front Mol Biosci 7, 586413 (2020).

38. J. G. Belasco, J. T. Beatty, C. W. Adams, A. von Gabain, S. N. Cohen, Differential expression of photosynthesis genes in R. capsulata results from segmental differences in stability within the polycistronic rxcA transcript. Cell 40, 171–181 (1985).

39. M. J. Weickert, S. Adhya, The galactose regulon of Escherichia coli. Mol Microbiol 10, 245–251 (1993).

40. S. Adhya, W. Miller, Modulation of the two promoters of the galactose operon of Escherichia coli. Nature 279, 492–494 (1979).

41. D. E. Lewis, S. Adhya, Molecular Mechanisms of Transcription Initiation at gal Promoters and their Multi-Level Regulation by GalR, CRP and DNA Loop. Biomolecules 5, 2782–2807 (2015).

42. S. Semsey, K. Virnik, S. Adhya, Three-stage regulation of the amphibolic gal operon: from repressosome to GalR-free DNA. J Mol Biol 358, 355–363 (2006).

43. K. A. Datsenko, B. L. Wanner, One-step inactivation of chromosomal genes in Escherichia coli K-12 using PCR products. Proc Natl Acad Sci U S A 97, 6640–6645 (2000).

44. X. Wang, H. J. Jeon, M. P. A. N J. He, H. M. Lim, Visualization of RNA 3’ ends in Escherichia coli Using 3’ RACE Combined with Primer Extension. Bio Protoc 8, e2752 (2018).

45. M. P. A. N H. M. Lim, An in vitro Assay of mRNA 3’ end Using the E. coli Cell-free Expression System. Bio Protoc 12, e4333 (2022).

46. S. C. Ji et al., In vivo transcription dynamics of the galactose operon: a study on the promoter transition from P1 to P2 at onset of stationary phase. PLoS One 6, e17646 (2011).

47. X. Wang, M. P. A. N H. J. Jeon, J. He, H. M. Lim, Identification of a Rho-Dependent Termination Site In Vivo Using Synthetic Small RNA. Microbiol Spectr 11, e0395022 (2023).

