## Supplementary material for "Rho-dependent termination and RNase E-mediated cleavage: Dual pathways for RNA 3’ end processing in polycistronic mRNA"

**Short title:** RNA 3’ end formation via Rho and RNase E

**This PDF file includes:**

Figures S1 to 4
Table S1

**Supplementary figures**

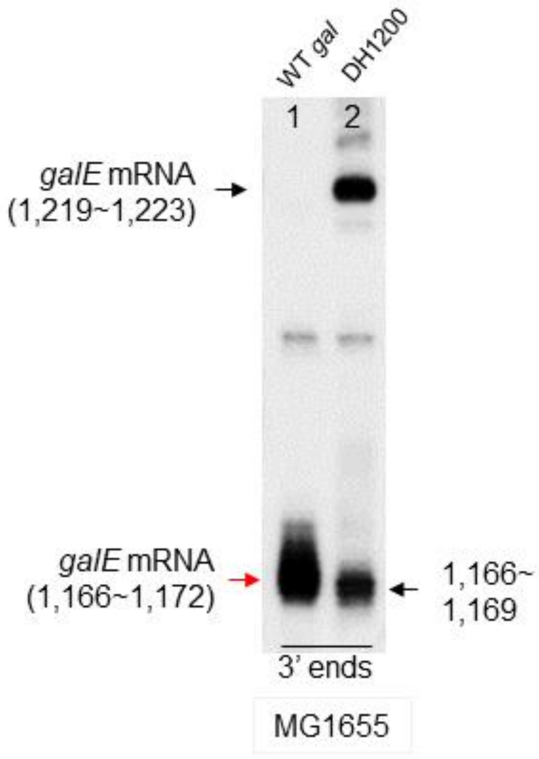


**Fig S1. 3′ RACE assay of *galE* mRNA in the *DH1200* mutant with an additional E-hairpin at 1,200 (lane 2).** The E-hairpin (42 nucleotides; 1,137 to 1,178) at position 1,200 in the WT *gal* operon was inserted to generate the double-hairpin *gal* mutant DH1200 (1). Related to Figure. 2.

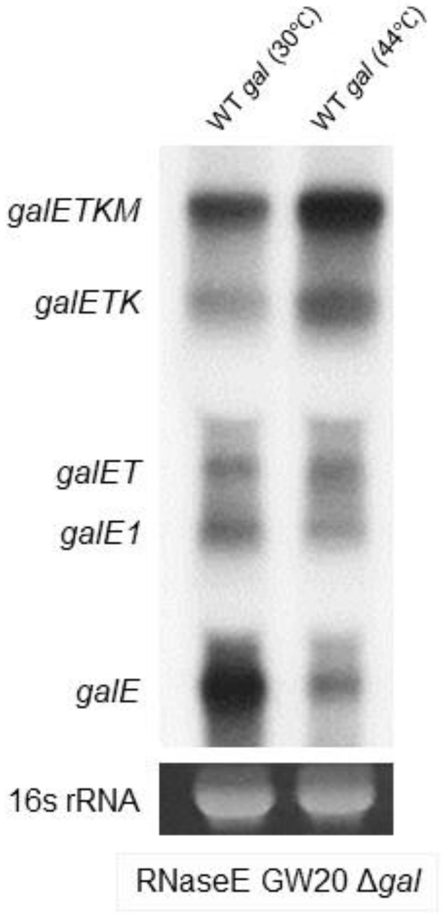


**Fig S2. RNase E-mediated endo-nucleolytic cleavage processing is the source of the 3’ ends of pre-*galE*.** Northern blotting was used to analyze the total RNA from the cell cultures (see Materials and Methods) of *gal* mRNAs in GW20Δ*gal* (temperature-sensitive RNase E mutant) cells. Cells were cultured at both permissive (30°C) and non-permissive (44°C) temperatures for analysis. Related to Figure. 3A.

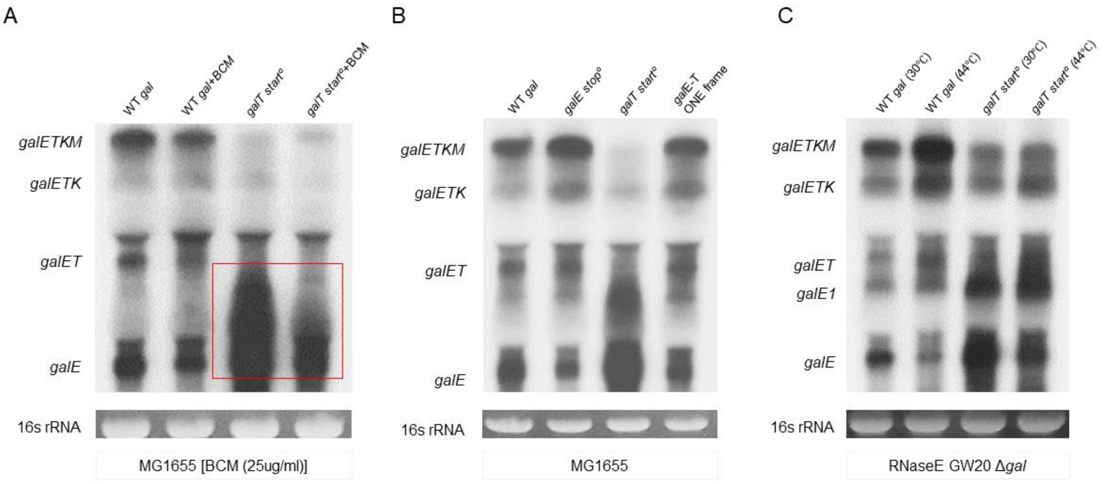


**Fig S3. Failure of *galT* translation initiation causes RDT. A)** Northern blot analysis of *gal* mRNAs in *galT* start$^{o}$ mutant with or without the Rho inhibitor bicyclomycin (BCM). To determine if *galE* mRNA is produced by RDT, we treated MG1655 cells at an OD600 of 0.6 in LB medium with 25µg/mL BCM concentration for 10 minutes. Northern blotting was used to analyze the total RNA from the cell cultures (see Materials and Methods). Related to Figure 5A. B) Northern blot analysis of *gal* mRNA levels in the *galE* stop$^{o}$, *galT* start$^{o}$, and *galE-T* ONE frame mutants. A slight increase of the other *gal* mRNAs longer than *galE* in the *galE stop$^{o}$* mutant could have resulted from the fact that more transcription might have gone downstream of the 3’ end of *galE*. C) Northern blot analysis of *gal* mRNA levels in the *galT* start$^{o}$ mutant in GW20Δ*gal* (temperature-sensitive RNase E mutant) cells. Cells were cultured at both permissive (30°C) and non-permissive (44°C) temperatures for analysis. Related to Figure. 5B.

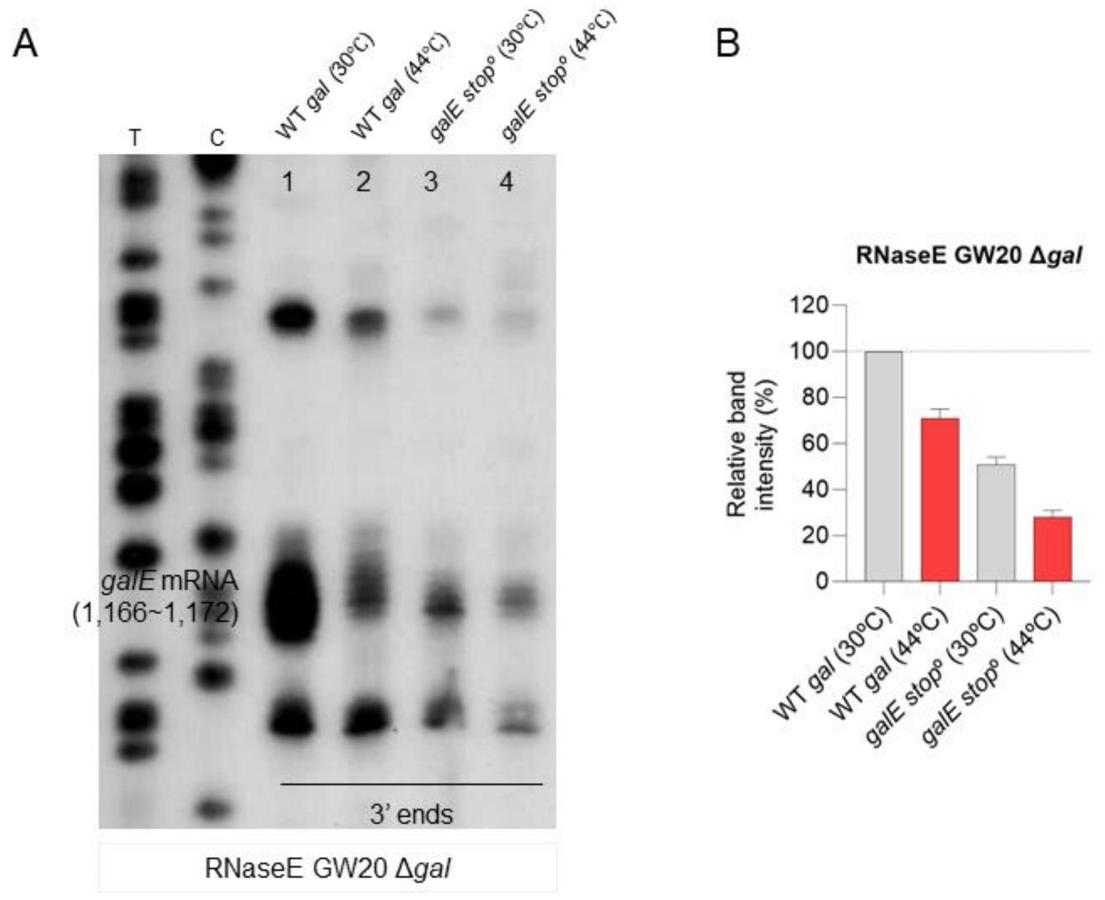


**Figure S4. Continuous translation without interruption by termination at the stop codon of *galT* could eliminate the break in transcription-translation coupling.** The 3'RACE assay compares the WT and *galE* stop° mutant in GW20Δ*gal* (temperature-sensitive RNase E mutant) cells, where the whole *gal* operon is deleted from the chromosome. Cells were cultured at both permissive (30 °C) and non-permissive (44 °C) temperatures for analysis. B) Relative band intensity for the 3'RACE assay from (A).

**Supplementary table**

Supplementary Table 1. Primers used in this study and usage.

| **Primer name** | **Primer sequences (5′→3′)** | **Usage** |
|---|---|---|
| *galBamHI-R* | CCGGATCCGGTGATTTGAACAATATGAG | Cloning |
| *galHindIII-F* | CACCGTTTATGGCGATCAGCCC | |
| *galMluI-R* | GCGTTTTCAGTCAGTATATGACG | |
| *EHMM2-F* | CGGCCCCTGGCAGGGGGCGCAGGAAA | Site-directed mutagenesis |
| *EHMM5-F* | CGCGGCCTGGCAGGGGGCGCAGGAAA | |
| *ENO-stop-R* | GGTCGTTCCTTTATCCGGATATC | |
| *TNO-start-F* | TAAGGAACGACCAAAACGCAA | |
| *ET-one frame* | AAAGGAACGACCAAAACGCAA | |
| *DH1200aatII-F* | GTTACCTGCGCACGATCCAGACGTC | |
| *DH1200aatII-R* | ACATTACCTGCGCAGAGGAAGACGTC | |
| *850 MluI-R* | CCATTACGCGTATGGTATGAAATAACC | |
| *850 PstI-R* | CAAATTCACGCCTGCAGGCGCCTGATT | |
| *E1-F* | ATGAGAGTTCTGGTTACCGGTGGTAG | E-probe generation for northern blot |
| *E1-R* | TGGGCTTTTTTGCAGATCGGTGAGGA | |
| *3RP* | AGCATGCGGCCGCTAAGAAC | 3' RACE – RT and PCR primers |
| *3' RNA Oligo* | UUCACUGUUCUUAGCGGCCGCAUGCU | |
| *E3-F* | CATCGCCCAGGTTGCTGTAG | |
| *NewT6_1123-F* | TTTCACCGCACCGCGCTAA | 3' RACE extension primer |
| *T1, T-ext1,T6-F* | ATGACGCAATTTAATCCCGTTGATCATC | |
| *5s-F* | GAGAGTAGGGAACTGCCA | 5′ RACE PCR primer |
| *K2-R* | AGCCTACAAACTGGTTTTCTGCTTCC | |
| *M2-1-R* | CATCTGAACTCAGGGCAAACA | |
| *EText-R* | AGAATCCATTGCCCGGTGAG | 5`RACE extension primer |
| *TKext-R* | ATGGTGTGAGTGGCAGGGTA | |
| *KMext-R* | TGCCAGTGCGGGAGTTTCGT | |

**References**


1. Jeon HJ, N MPA, Lee Y, Lim HM. 2022. Failure of Translation Initiation of the Next Gene Decouples Transcription at Intercistronic Sites and the Resultant mRNA Generation. *mBio* 13:e0128722.
2. Wang X, N MPA, Jeon HJ, Lee Y, He J, Adhya S, Lim HM. 2019. Processing generates 3' ends of RNA masking transcription termination events in prokaryotes. *Proc Natl Acad Sci U S A* 116:4440-4445.
3. Wang X, Ji SC, Yun SH, Jeon HJ, Kim SW, Lim HM. 2014. Expression of each cistron in the *gal* operon can be regulated by transcription termination and generation of a *galk*-specific mRNA, mK2. *J Bacteriol* 196:2598-606.
